# AlphaFold predicted structure of the Hsp90-like domains of the neurodegeneration linked protein sacsin reveals key residues for ATPase activity

**DOI:** 10.1101/2022.10.18.512535

**Authors:** Laura Perna, Lisa E. L. Romano, Chrisostomos Prodromou, J. Paul Chapple

## Abstract

The ataxia-linked protein sacsin has three regions of partial homology to Hsp90’s N-terminal ATP binding domain. Although a crystal structure for the Hsp90-like domain of sacsin has been reported the precise molecular interactions required for ATP-binding and hydrolysis are unclear. To better understand how sacsin may function as an ATPase we utilized an AlphaFold predicted structure of its Hsp90-like domain. Superimposition onto Hsp90, and other modelling approaches, have resulted in novel insights into sacsin’s structure. These encompass identification of residues within the sacsin Hsp90-like domains that are required for ATP binding and hydrolysis, including the catalytic arginine residues equivalent to that of the Hsp90 middle domain. Importantly, our analysis allows comparison of the Hsp90 middle domain with corresponding sacsin regions and has identified that sacsin has a shorter lid segment than the N-terminal domain of Hsp90. We also speculate, from a structural viewpoint, why ATP competitive inhibitors of Hsp90 do not appear to affect sacsin. Together our analysis supports the hypothesis that sacsin’s function is ATP-driven and would be consistent with it having a role as a molecular chaperone. We propose that the SR1 regions of sacsin be renamed as HSP-NRD (Hsp90 N-Terminal Repeat Domain; residues 84-324) and the fragment immediately after as HSP-MRD (Hsp90 Middle Repeat Domain; residues 325-518).

## Introduction

Mutations which lead to loss of function of the protein sacsin cause the neurodegenerative disorder Autosomal Recessive Spastic Ataxia of Charlevoix Saguenay or ARSACS [1-3]. Although a very rare disease, ARSACS is thought to be the second most common form of autosomal recessive cerebellar ataxia after Friedrich’s ataxia [2]. It normally manifests in childhood and is characterised by progressive cerebellar ataxia, peripheral neuropathy, and spasticity.

Sacsin is an extremely large (4579 amino acid) multidomain protein that is conserved through vertebrate evolution [4], with potential orthologs identified in multiple eucaryotes including *Acropora corals* [5]. There is no overall structural similarity between sacsin and other proteins, however it does contain conserved domains (Figure 1). Specifically, from the N- to C-terminus sacsin incorporates; (i) a ubiquitin-like domain (UBL) that interacts with the 19S cap of the 26S proteasome and mediates protein degradation [6,7]; (ii) three supra domains known as sacsin internal repeats (SIRPT), that can be further divided into smaller sub-repeats known as SR1, SR2, SR3 and SX (SIRPT2 lacks the SRX repeat), with each SR1 containing a region of homology to the Hsp90 N-terminal ATPase domain [4,8]; (iii) a xeroderma pigmentosum complementation group C binding (XPCB) domain that interacts with the ubiquitin ligase and Angelman syndrome protein Ube3A [9,10]; (iv) a J-domain that binds and activates Hsp70 [6,8]; and, (v) a higher eukaryotes and prokaryotes nucleotide-binding domain (HEPN) that may promote sacsin dimerization [11,12].

**Figure 1.**
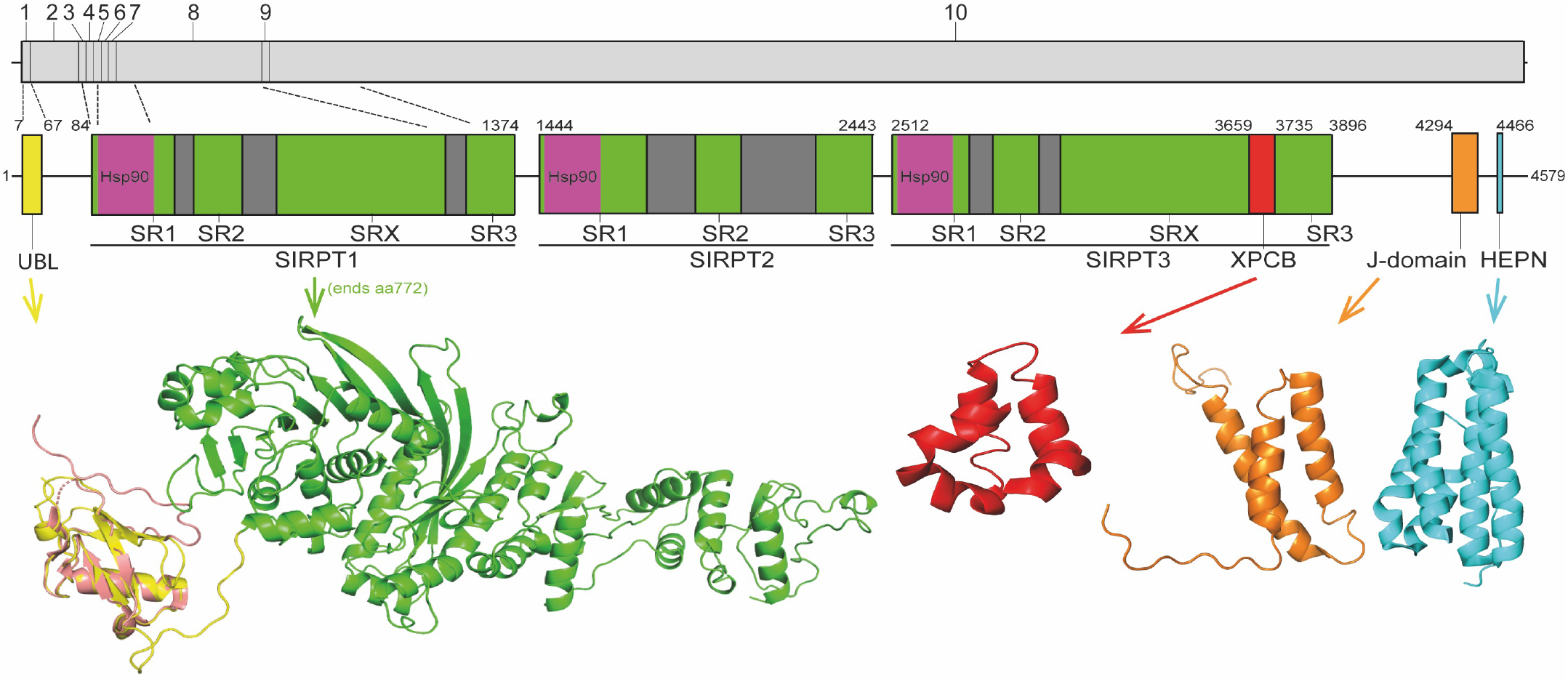
Schematic representation of sacsin domain structure. Regions of sacsin with homology to domains of other proteins are indicated (numbers correspond to amino acid position) with domains where structure has been solved or modelled shown (UBL, Hsp90, XPCB, J-domain and HEPN). The internal repeats of sacsin are also indicated and the structure of the sacsin transcript is also shown (with exons numbered and dotted lines indicating exons encoding different regions of sacsin).

This domain structure links sacsin to both the ubiquitin proteasome system and molecular chaperones, suggesting a potential function in protein quality control systems. Although there is some evidence supporting this [6,13], the precise role of sasin is unknown and it is unclear why its loss results in a complex cellular phenotype that includes mitochondrial dysfunction [11,14,15], altered intermediate filament cytoskeleton organisation [13,16,17], altered microtubule dynamics [18] and disrupted intracellular trafficking [19].

One possibility is that sacsin functions directly as a molecular chaperone and if this is the case its three regions of homology to Hsp90 (in the SR sub-repeats of SIRPT1, 2 and 3) could indicate its action, like Hsp90, is driven by ATP binding and hydrolysis. Currently, sequence alignment and crystallographic analysis have shown that the sacsin SR1 has structural similarity to the Bergerat protein fold of Hsp90, which forms a nucleotide binding pocket from a sandwich of four β-sheets and two α-helices [7]. Sacsin’s SR1 Bergerat fold is not entirely structurally conserved with that of Hsp90 proteins but does show conservation of amino acids important for ATP binding and hydrolysis. However, it is not clear whether SR1 has a segment of structure that could act as an ATP-binding lid, which may be consistent with recombinant SIRT1-SR1 having low levels of ATPase activity in a steady state assay [7]. Also, this conflicts another study where a larger domain of mouse sacsin, from the N-terminus to start of SIRT2, was generated as a recombinant protein and demonstrated to have ATPase activity equivalent to yeast Hsp90 [8]. These data may be consistent with a region of sacsin outside of the SR1 domain contributing to its ATPase activity. It is also interesting to note that key residues in the SR1 putative nucleotide binding pocket are completely conserved between the SIRT domains of sacsin, suggesting they are important for function [7]. Moreover, there is evidence for the ATPase function of the SR1 being important for sacsin function, as it has been shown that the ARSACS mutation D169Y suppresses sacsin activity [8]. Importantly, Hsp90 possesses a catalytic-loop arginine, found on its middle-domain that is required to complete the ATPase unit of Hsp90 and thus allow hydrolysis of ATP, yet at first sight sacsin appears to have no obvious domains equivalent to the middle domain of Hsp90.

To resolve this controversy and better understand the molecular function of sacsin we have exploited recent advances in protein structural modelling to identify key residues within sacsin that could mediate ATP binding and catalysis.

## Materials and Methods

### Superimposition of Hsp90 and sacsin structures and sequence alignment

The N-terminal domain of Hsp90 was superimposed with the SPRPT1-SR1 domain of sacsin from the AlphaFold structure (UniProt entry A0A804HIU0) using pymol (The PyMOL Molecular Graphics System, Version 1.2r3pre, Schrödinger, LLC). Once superimposed, the peptide sequences between Hsp90 and sacsin were aligned and residues conserved in Hsp90 for ATP binding and hydrolysis were compared to residues in the sacsin sequence.

### Superimposition of the middle domain of Hsp90 and sacsin

The catalytic arginine of sacsin was identified following the superimposition of the N-terminal domain of Hsp90 onto the Hsp90-NRD of the AlphaFold structure of sacsin, by using PyMol (The PyMOL Molecular Graphics System, Version 1.2r3pre, Schrödinger, LLC). The best superimposition of the middle domain of Hsp90 (residues 262 to 408) and sacsin (residues 325 to 518), was obtained by orientating the Hsp90 middle domain so that the Hsp90 middle domain β-sheet matched with that of sacsin. We maintained the antiparallel and parallel nature (single pair of strands) of the β-strands that formed each of the corresponding β-sheets and allowed the longest helix of both the Hsp90 middle domain and the equivalent sacsin segment to be approximately aligned. Using the structural alignment, peptide sequences on either side of the conserved arginines of sacsin were aligned.

### Treatment of SH-SY5Y wells with Hsp90 inhibitors and immunoblotting for sacsin

SH-SY5Y cells were purchased from the European Collection of Authenticated Cell Culture and grown in DMEM-F12 supplemented with 10% heat-inactivated fetal bovine serum and antibiotics (50U/ml penicillin and 50μg/ml streptomycin). Cells at approximately 80% confluence were treated with the Hsp90 inhibitors geldanamycin or tanespimycin/17-AAG at final concentrations of 5, 10 or 15µM, or a vehicle only control for 24 hours. Total cell lysates were then harvested in RIPA Buffer (150 mM NaCl, 1.0% IGEPAL CA-630, 0.5% sodium deoxycholate, 0.1% SDS, and 50 mM Tris, pH 8.0), supplemented with protease (ThermoFisher, UK) and phosphatase inhibitors (Roche, Burgess Hill, UK), and Western blotted for sacsin (using anti-sacsin antibody ab181190 [Abcam] at a titre of 1:1500) the Hsp90 client AKT (ab 9272S [Cell Signalling] 1:1000) and β-actin (ab 8226 [Abcam] 1:10000) as a loading control.

## Results

### The AlphaFold predicted sacsin SR domains are structurally consistent with the ATPase constraints of the Begarat fold

Recently the structure of a region of human Sacsin (amino acids 1 to 177) has been determined using AlphaFold [20,21]. Using the deposited structure, Uniprot A0A804HIU0, we have investigated whether the SR1 domains of the SIRPT regions of sacsin would be structurally consistent with the ATPase constraints of the Begarat fold. The crystal structure of the SR1 domain of sacsin (PDB 5V44) is essentially the same as that for the AlphaFold prediction, except that the crystal structure lacks details for the lid. In another deposited structure, PDB 5V46, the lid is almost intact, but not in a closed conformation and thus does not superimpose with the AlphaFold predicted structure. Using PyMOL [22] to superimpose the N-terminal domain of AMPPNP bound Hsp90 (PDB 2CG9) with the SIRPT1-SR1 domain of sacsin, we were able to confirm that sacsin contains a Begarat fold, as reported earlier (Figure 2A) [7]. We also confirm that the lid segment of the AlphaFold sacsin structure appears to be in a closed position, similar to that of Hsp90 but is apparently shorter in overall length.

**Figure 2.**
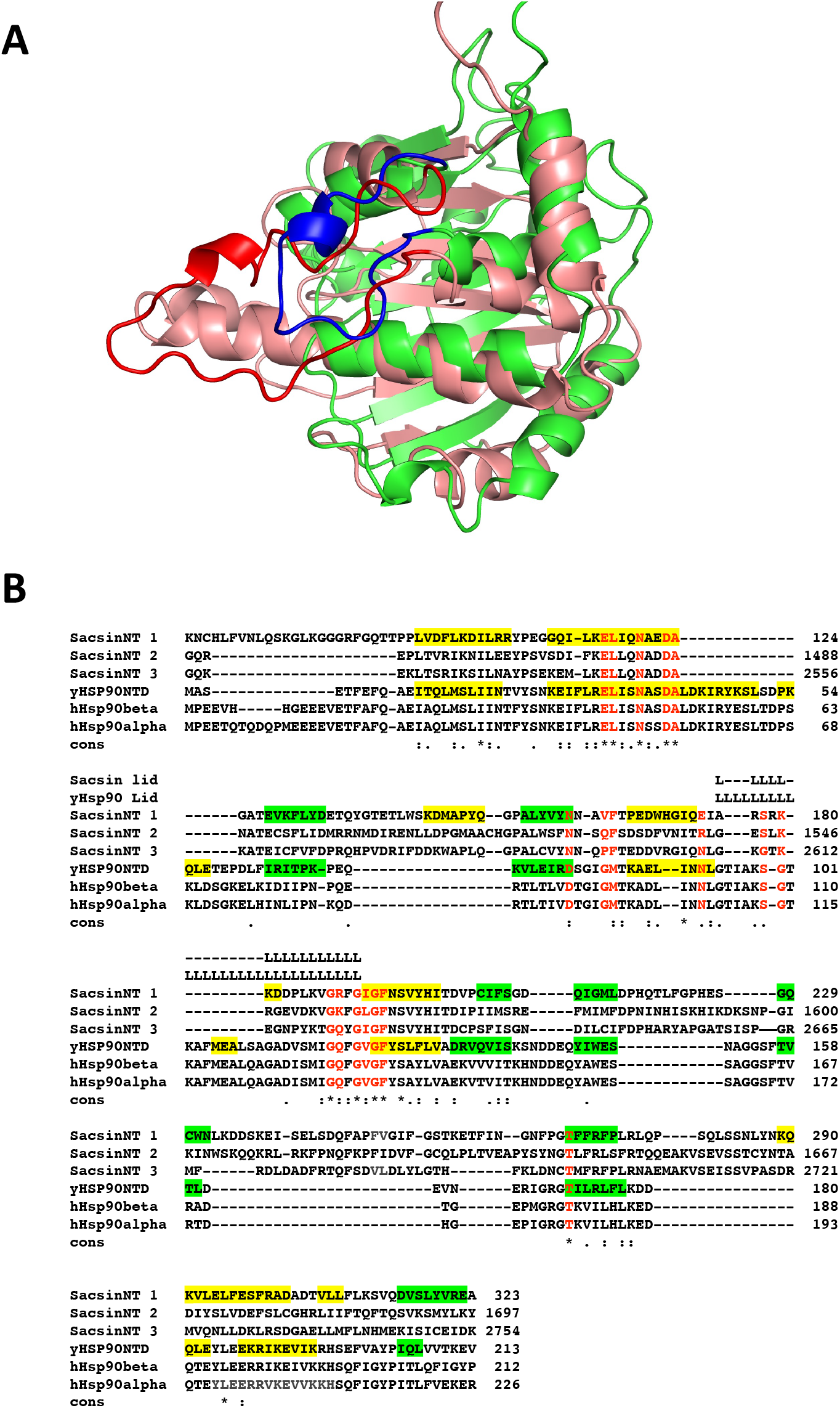
Superimposition of the Sacsin SR1 and Hsp90 N-terminal domains. A, The N-terminal domain of Hsp90 (salmon with red lid, PDB 2CG9) was superimposed onto the SR1 domain of the AlphaFold UniProt entry A0A804HIU0 (green with blue lid). B, Sequence alignment of the N-terminal domains of yeast (yHsp90NTD) and human Hsp90 (hHsp90 α and β) with the three SR-domains of sacsin (SacsinNT 1 to 3). Green highlights, β-strand and yellow, α-helix. Cons, conservation, (.), weakly conserved residue position; (:) strongly conserved residue position and (*), invariant residue position.

### Structure based alignment identifies residues required for ATP binding in each of sacsin SR domains

Using the structurally aligned superimposition of Hsp90 and sacsin, we next compared the amino acid residues of Hsp90 involved in binding and hydrolysis of ATP with the peptide sequence of sacsin (Figure 2B). Using this alignment allowed us to accurately determine which sacsin residues correspond to ATP binding residues of Hsp90. The structurally based alignment shows that many of the residues required for ATP binding and hydrolysis are conserved in sacsin (Figure 3). The catalytic Glu 33 (sacsin Glu 116) as well as other residues involved in binding of ATP are invariant. These include Leu 34 (sacsin Leu 117-where a single sacsin position is given in the following description, the residues are conserved in all three SIRPT-SR1 domains and only differences to SIRPT1-SR1 residues are reported from here on), Asn 37 (sacsin Asn 120), Asp 40 (sacsin Asp 123), Ala 41 (sacsin Ala 124), Gly 118 (sacsin 188), Gly 121 (sacsin Gly 191), Gly 123 (sacsin Gly 193), Phe 124 (sacsin Phe 194) and Thr 171 (sacsin Thr 269). Conserved residues include Asp 79 (sacsin Asn 160), Met 84, (sacsin Phe 164), Asn 92 (sacsin Gln 173 and Thr 1539 for the SR2 domain) and Ser 99 (sacsin Ser 179 and Gly 210 for SIRPT3-SR1). Previously, it was claimed that Sacsin Phe 164 aligned with Hsp90 Gly 83 [7]. However, our analysis suggests that Phe 164 replaces Hsp90 Met 84, allowing Phe 164 to perhaps pi-stack with the adenine ring of ATP. Consequently, Hsp90 Gly 83 aligns with SIRPT1-SR1 Val 163 (Q1529 for SIRPT2-SR1 and P2595 for SIRPT3-SR1) Val163, Q1529 and Pro 2595 could all mimic the main-chain interactions formed by Gly 83 via a water molecule to Asp 79 and to a nitrogen atom of the adenine ring of ATP. Thus, the side chains of valine, glutamine and proline would all point away from the bound ATP and would not therefore interfere with its binding.

**Figure 3.**
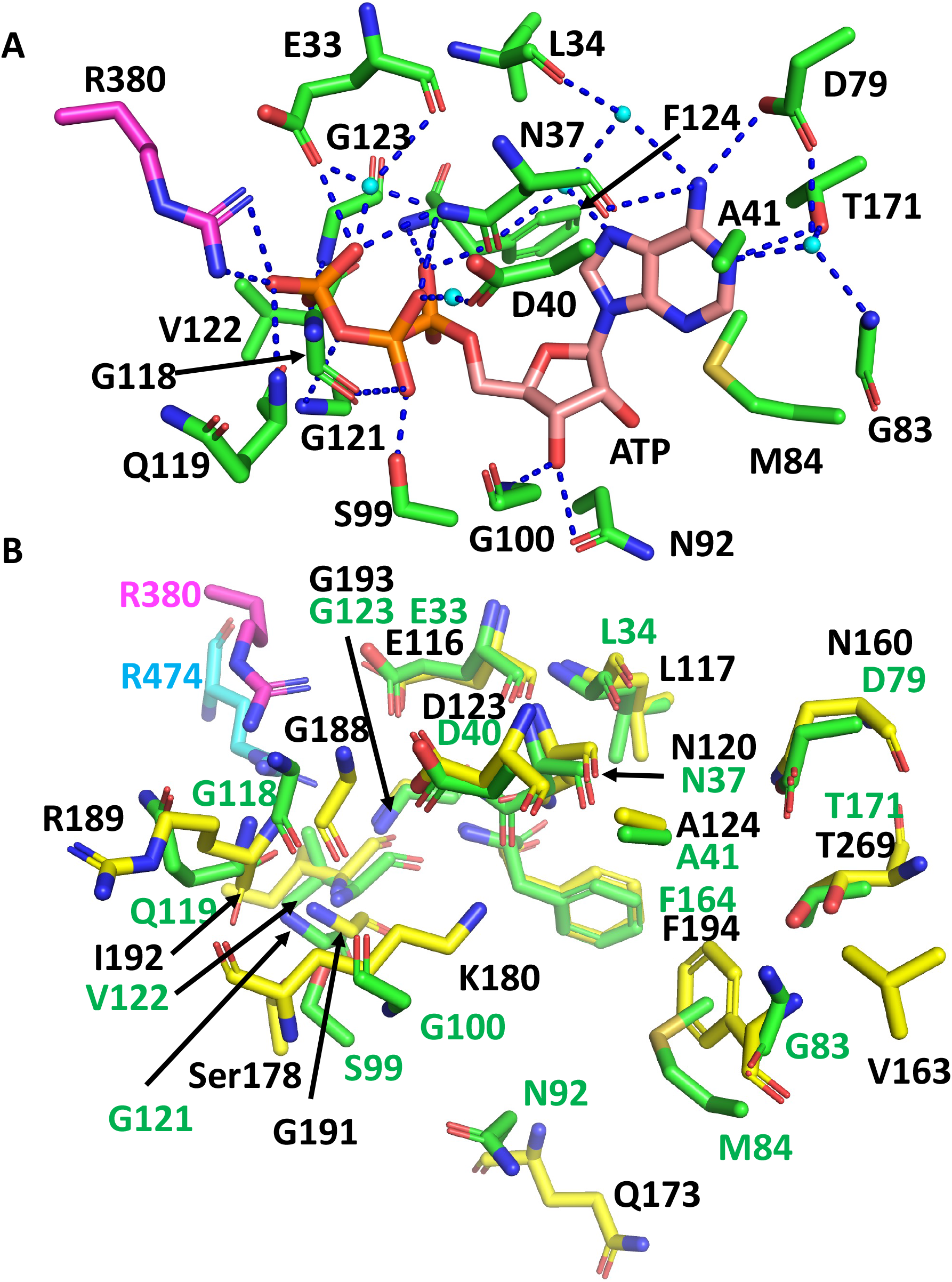
Comparison of the residues required for ATP binding and hydrolysis by Hsp90 and Sacsin. A, PyMOL cartoon of the yeast Hsp90 residues involved in the binding and hydrolysis of ATP. Residues are in stick conformation. The catalytic Arg 380 of Hsp90 is coloured in magenta. Water molecules are shown as cyan coloured spheres and hydrogen bond interactions as dashed blue lines. B, PyMol cartoon showing the superimposition of Hsp90 residues involved in the binding and hydrolysis of ATP and the corresponding sacsin residues. Green stick residues with green numbering, Hsp90 residues with the catalytic Arg 380 in magenta. Yellow stick residues with black numbering, sacsin residues with the catalytic Arg 474 shown in cyan.

Other main chain contacts between Hsp90 and ATP include Val 122 that would allow substitutions in sacsin (Ile 192 and Leu 1558 for SIRPT3-SR1) and Gly 100 (sacsin Lys 180) which is replaced by lysine in sacsin. Finally, the main chain of Gln 119 contacts ATP and this interaction could be maintained with the substitutions seen in sacsin (Arg 1555 and Lys 2621 of the SIRPT2-SR1 domain).

### The lid structure of the ATP binding pocket is shorter in sacsin than Hsp90

On closer inspection, the greatest variability in ATP binding residues between Hsp90 and sacsin are those that occur on the lid region of the SR domains. We note that the lid structure of sacsin is significantly shorter than that of Hsp90 (Figure 2A). Sacsin residues that do not superimpose well with corresponding ATP binding residues of Hsp90 in the structure alignments include Ser 178 (Hsp90 Ser 99), Lys 180 (Hsp90 Gly 100), Gly 188 (Hsp90 Gly 118) and Arg 189 (Hsp90 Gln 119) (Figure 3B). The substitution of Gln 119 with Sacsin Arg 189 can maintain the main chain interaction with ATP and the side chains of these amino acids are pointing away from the bound ATP. However, in the case of the substitution of Gly 100 with sacsin Lys 180, this at first sight appears to cause a clash with bound ATP. However, this residue position is solvent exposed, which might allow the side chain of Lys 180 to adopt a conformation that allows unhindered ATP binding, while maintaining a main chain contact with ATP. Alternatively, the side chain of Lys 180 might adopt one of a number of other possible conformations that could perhaps allow it to interact with sacsin Asn 120 and Asp 123 or even with the phosphate or 2’ and or 3’ hydroxyls of the ribose sugar of the bound ATP. However, the fact that in Hsp90 this position is an invariant glycine (Gly 100), does mean that a crystal structure is required to establish the exact consequences of this lysine substitution on the ATPase activity of these sacsin domains. Nonetheless, the misalignment of specific lid residues between Hsp90 and sacsin means that it is likely that the model in this region of the AlphaFold structure either needs further refining or that the local structure of the lid is restructured upon ATP binding.

### The equivalent residues in sacsin to the catalytic arginine in the middle domain of Hsp90 are predicted to be Arg 474, Arg 1839 and Arg 2893

With the N-terminal domain of Hsp90 correctly superimposed onto sacsin SIRPT-SR1, we were able to locate the catalytic arginine required for ATPase activity in the downstream (SIRPT1) region of sacsin, as observed in the Hsp90’s middle domain [23]. We were able to identify Arg 474 of sacsin being orientated close enough to the superimposed Hsp90 bound ATP molecule and able to form similar contacts to ATP as seen with the catalytic Arg 380 of yeast Hsp90 (Figure 4A). We therefore propose that Arg 474, Arg 1839 and Arg 2893, which are conserved residue positions in SIRPT1-3 regions of sacsin, are the catalytic arginine residues required by the SR1 domains for catalysis (Figure 4B). The alignment of the arginine catalytic loop, and the structural elements on either side of this loop, are presented in Figure 3B, which shows little peptide sequence conservation with yeast Hsp90. The conservation of Arg 474, 1839 and 2893, suggests that sacsin would be an active ATPase. Furthermore, the R474C mutation in sacsin has been reported to score as highly as truncated forms of sacsin on the SPAX Scoring System, suggesting that R474C mutant is non-functional [19]. This provides further support that sacsin is an active ATPase protein.

**Figure 4.**
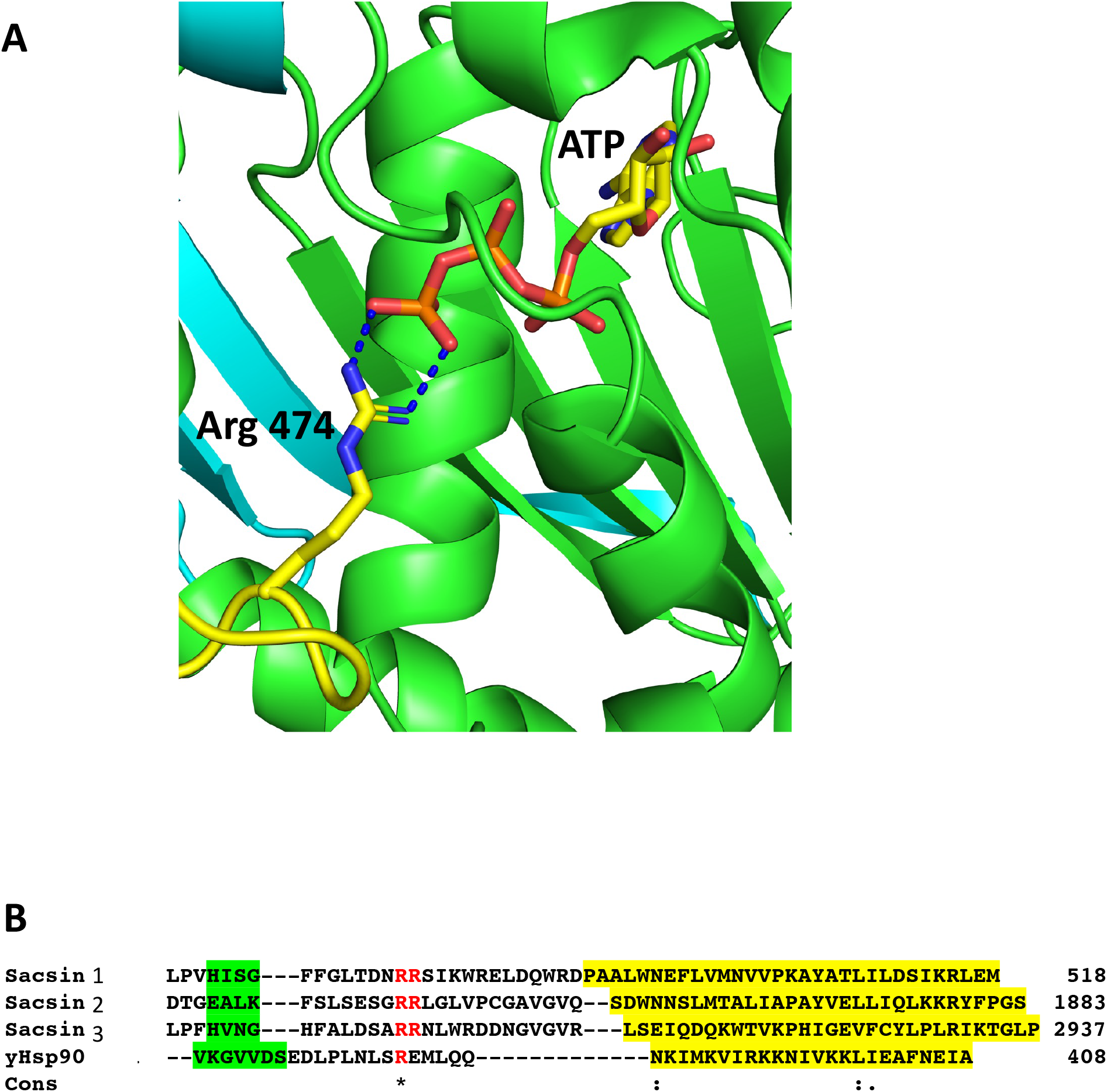
The catalytic Arg 474 of sacsin. A, PyMol cartoon of the potential interaction of Arg 474 with bound ATP in the SR domain (green). B, Alignment of the catalytic loop regions, containing the conserved arginine residues of SR1, 2 and 3 (Sacsin 1, 2 and 3), and the adjoining structural elements with the same regions as for yeast Hsp90 (yHsp90). Green highlight, β-strand and yellow highlight, α-helix. Cons, conservation, (.), weakly conserved residue position; (:), strongly conserved residue position and (*), invariant residue position.

### Structural similarity exists between the middle domain of Hsp90 and sacsin SIPRT domains

In order to obtain the best superimposition of the middle domain of Hsp90 with the equivalent region of sacsin, an orientation of the Hsp90 middle domain (residues 262 to 444 used) was required. The middle domain of Hsp90 contains a 7-stranded β-sheet, whereas sacsin contains a continuous 13-stranded β-sheet that runs from the SR1 domain and into the downstream SIRPT1 region, of which the last 5 β-strands appear to form the central core of the domain (Figure 5A). To get the best superimposition of the middle domain of Hsp90 (residues 262 to 408) with sacsin (residues 325 to 518), we orientated the Hsp90 middle domain β-sheet such that we could match it with that of sacsin (Figure 5B). The best alignment maintained the antiparallel and parallel nature (single pair of strands) of the β-strands that formed each β-sheet and allowed the longest helix of both the Hsp90 middle domain and the equivalent sacsin segment to be approximately aligned (Figure 5C). From this superimposition, a central core of structure that could be considered similar was identified (Figure 5C, D). Collectively this included the superimposed β-strands and the long helix of these domains. Most importantly, the structural elements on either side of the arginine catalytic loop, which consist of one of the superimposed β-strands and the following superimposed long helix of these domains were matched well. This suggested a similar sub-structure (Figure 5C) in what otherwise appears to be an unrelated fold at first sight (Figure 5B). The topology of the Hsp90 middle domain and the corresponding sacsin region is shown in Figure 5E. Consequently, we identify sacsin residues 325 to 518 as representing a Hsp90-like middle domain, which corresponds to a fragment of the Hsp90 middle domain (residues 262 to 408). We propose that the SR1 regions of sacsin be renamed as the HSP-NRD (Hsp90 N-terminal Repeat Domain; residues 84-324) and the fragment immediately downstream as HSP-MRD (Hsp90 Middle Repeat Domain; residues 325-518). Residues immediately after position 518 of sacsin do form α-helices, as seen in the Hsp90 middle domain, but our alignments between sacsin and Hsp90 of these secondary structural elements did not superimpose well.

**Figure 5.**
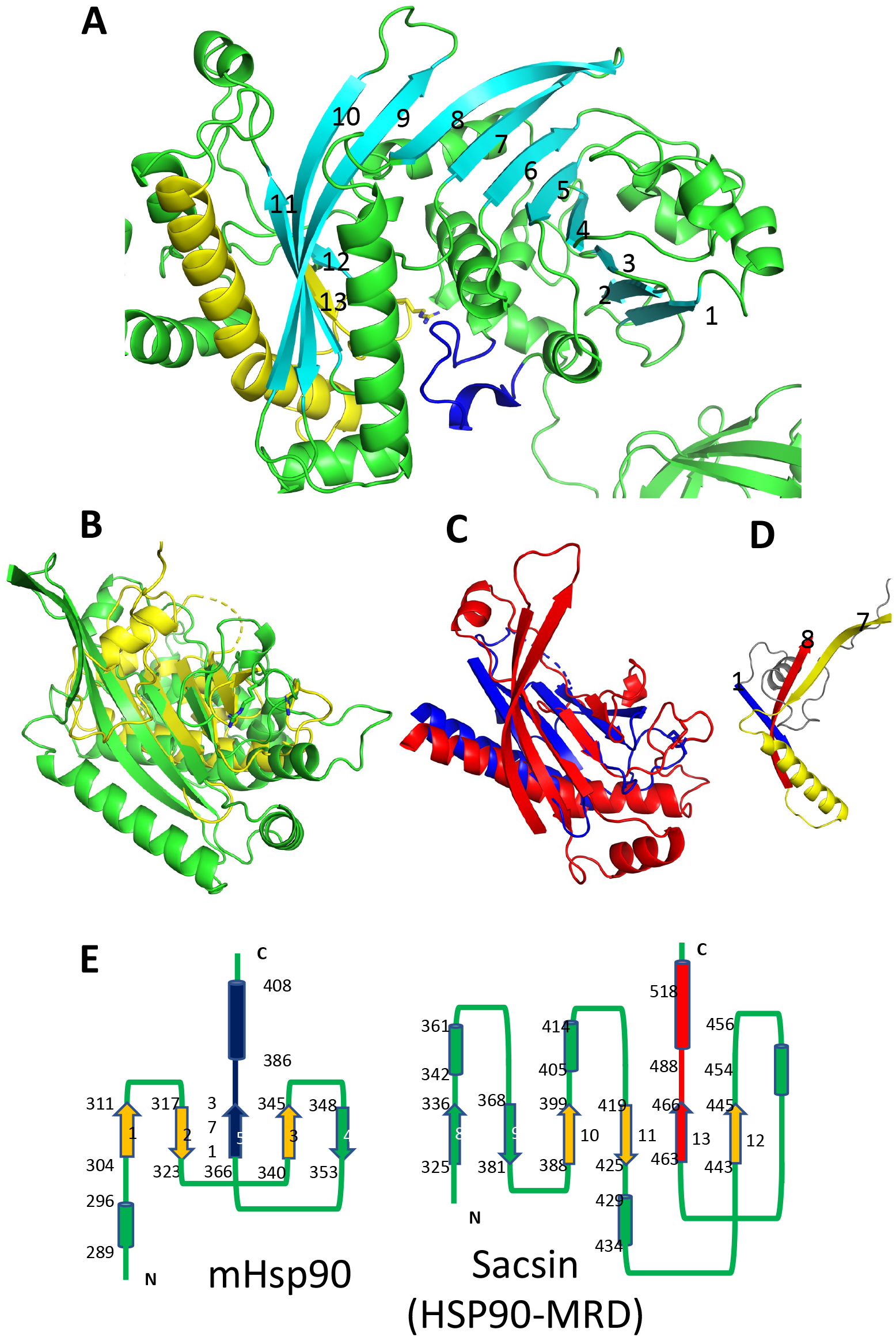
Comparison of the Hsp90 middle domain with corresponding sacsin regions. A, PyMol cartoon of the SR1 domain and immediate downstream region of sacin (green and cyan). A continues 13 stranded β-sheet runs from the SR1 domain and into the adjacent domain of sacsin. Cyan, β-strands, yellow, the β-strand and helix flanking the catalytic loop (yellow). Blue, the lid segment of the SR1 domain of sacsin. B, PyMol cartoon showing the superimposition of the Hsp90 middle domain (yellow, residues 262-444) and the corresponding sacsin region (green, residues 325-518). C, The central core structural elements of the middle domain of Hsp90 (blue) and the corresponding region of sacsin (red) show the main elements that superimpose. D, PyMol cartoon showing the N-terminal structural elements of the Hsp90 middle domain (blue and grey) and the corresponding elements of sacsin (red and yellow), showing that these structural elements do not superimpose. Strand 7 and 8, sacsin SR1 β-strands and (1), Hsp90 middle domain β-strand. E, Topology diagrams for the middle domain of Hsp90 (left panel) and the corresponding region of sacsin (right panel, HSP-Middle Repeat Domain (HSP-MRD)). Cylinders, α-helix, arrows, β-strand and lines are connections between the structural elements. The start and end residue numbers of each structural element are shown, as are the β-strand numbers. Blue and red, structural elements that represent the arginine catalytic loop and the flanking structural elements. Alignment of the red and blue β-strand and α-helix allows the superimposition of the orange β-strands.

Further analysis of the C-terminal domain of Hsp90 with topologically corresponding regions of sacsin, did not appear to show any structural homology (Figure S1). We therefore conclude that the SR1 domains of sacsin are homologues to the N-terminal domains of Hsp90, that the Hsp90 middle domain (residues 262 to 408) is structurally similar to a central core of the corresponding sacsin region, but the remaining section of Hsp90, including the C-terminal domain appears to be very different. None-the-less, it is clear that sacsin possesses a similar catalytic loop sub-structure that provides and orientates the catalytic arginine for catalysis.

### Hsp90 inhibitors are predicted not to bind sacsin because of steric hinderance from Asp 168

Hsp90 can be targeted by drugs that inhibit its ATPase activity through competitive binding of the nucleotide binding pocket (e.g., radicicol, geldanamycin and its analog tanespimycin/17-AAG) [24]. Given the homology between the Hsp90 ATP binding domain and the sacsin SR1 domains there is a possibility these inhibitors, or related drugs, may target sacsin. This has previously been investigated in an *in vitro* ATPase assay which saw no effect of geldanamycin or radicicol on activity of a region of mouse sacsin from the N-terminus of the protein to the beginning of the second SIRPT domain [8]. To provide a structural explanation of why these inhibitors appear not to affect sacsin we modelled superimposition of the N-terminal domain of Hsp90 containing geldanamycin and NVP-922 into sacsin’s ATP binding pocket (Figure 6). In Hsp90, the loop connecting β-strand 2 and the following α-helix consists of a total of 8 residues (positions 79 to 86). In contrast, sacsin has a shorter loop, consisting of just 6 residues (positions 160 to 166). The consequence of this is that the α-helix connected to this loop is drawn closer to bound ATP. While its effect on ATP binding appears negligible, it could result in clashes with geldanamycin and NVP-922 and in particular with the side chain of sacsin Asp 168 (Figure 6). This suggests that these inhibitors may not be able to bind directly to the ATPase site of sacsin.

**Figure 6.**
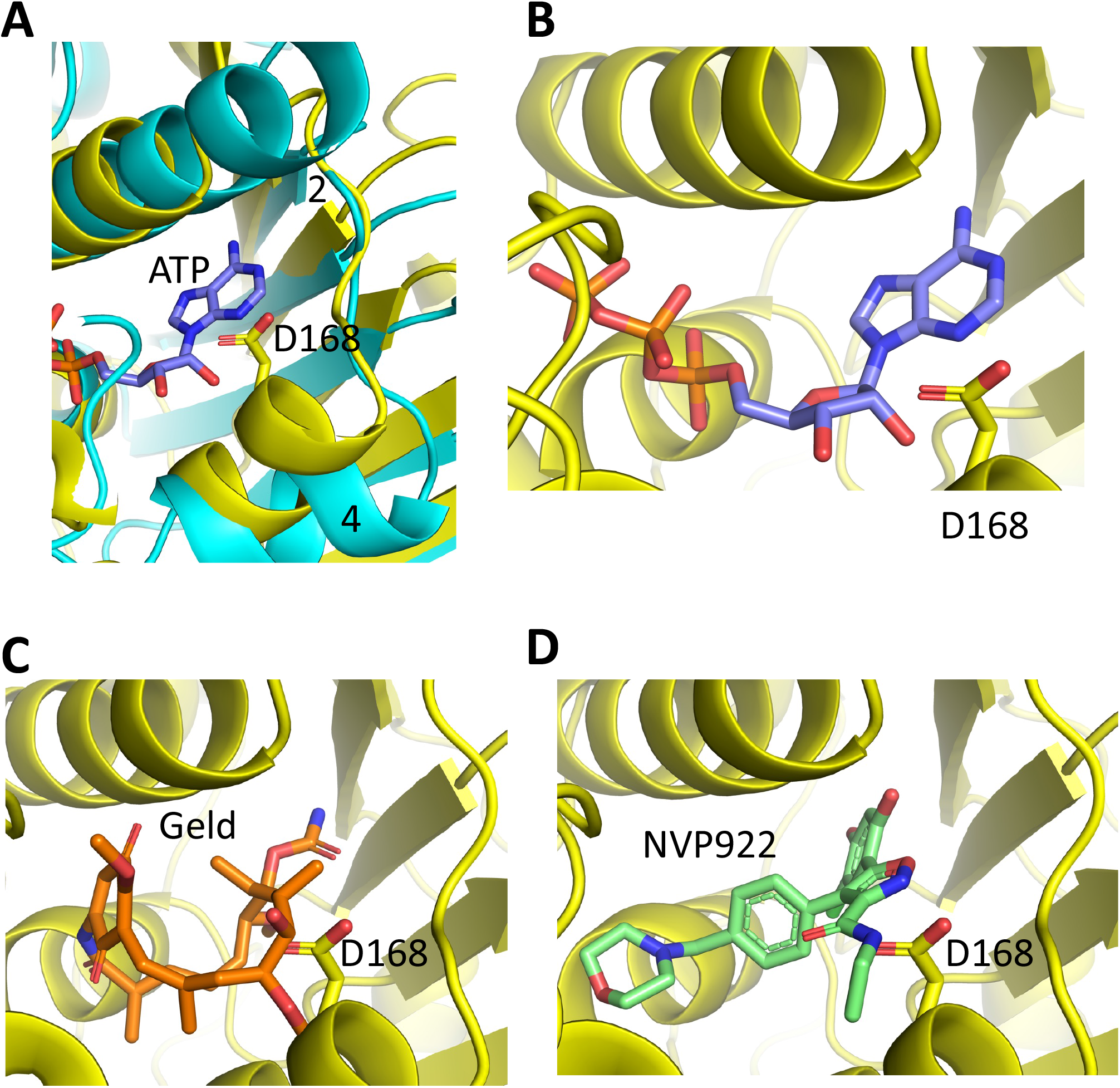
PyMol cartoons of Hsp90 N-terminal domain inhibitors superimposed into sacsins ATP binding pocket. A, α-Helix 4 of Hsp90 (cyan) is positioned to create a more open ATP-binding site. In contrast, the equivalent α-helix in sacsin (yellow) restricts the opening of the ATP-binding pocket. (2), β-strand 2 of Hsp90 is indicated. B, ATP docked into the ATP-binding site of sacsin, showing that it avoids any obvious clashes with the side chain of Asp 168. C, Geldanamycin docked into the ATP-binding site of sacsin, showing that it clashes with the side chain of Asp 168. D, NVP-922 docked into the ATP-binding site of sacsin, showing that it clashes with the side chain of Asp 168.

Our structural analysis indicates inhibitors that targeted the ATP binding site of Hsp90 are unlikely to directly affect sacsin. Yet, unexpectedly, we identified that treatment of the neuroblastoma-derived SH-SY5Y cell line with geldanamycin or 17-AAG leads to a reduction in cellular levels of sacsin (Figure S2). This may suggest that Hsp90 protein activity is required for normal expression or stability of sacsin.

## Discussion

Our analysis strongly supports that sacsin has three nucleotide binding domains with a similar structure to Hsp90 and that it is an active ATPase. This is despite differences between the lid of sacsin’s ATP binding pocket and the equivalent structure in Hsp90. More specifically, our alignments suggest that the lid segment of sacsin is about 12 residues shorter and could explain why the invariant Gly 100 of Hsp90 is replaced by Lys 180 in sacsin. ARSACS causing mutations are found in the predicted ATP binding regions of sacsin, including mutations that would be predicted to specifically inhibit ATPase activity (e.g., R474C), which supports the notion that sacsin requires ATP activity for its function.

The ATPase activity of Hsp90, and thus its function, is partly determined by dimerization mediated through the C-terminal domain [25-27]. Crystallographic structural analysis of sacsin’s C-terminal HEPN domain [28] also identified a dimer, suggesting the full-length protein might be dimeric. If this is the case, it is possible that dimerization of sacsin may influence the ATPase activity of its SR1 domains and ultimately its function. However, the AlphaFold structure shows that the lid segment of sacsin is packed against α-helix 1 and 2 including the loop following a-helix 2 of the HSP-MRD, which may indicate that direct dimerization with another Hsp90-like sacsin module is not required for ATPase activity. Interestingly, a-helix 1 and 2 of the HSP-MRD represents a structural element that does not superimpose with the Hsp90 middle domain. In contrast, Hsp90’s lid is mostly stabilised by the N-terminal domain from the adjacent protomer of the Hsp90 dimer. Thus, it appears that dimerization of the sacsin Hsp90-like domains may be very different, if it occurs at all, to that seen for Hsp90. However, assuming that dimerization of the Hsp90-like segments of sacsin is not required for its function, the question arises whether the mechanism of action between these chaperones is similar. Hsp90 has been seen to unfold clients such as kinases, with both protomers of the Hsp90 dimer involved in separating the N- and C-lobes of the kinase domain [29]. Alternatively, sacsin may use its individual Hsp90-like domains that are spatially separated, to achieve a similar effect on its clients, as seen with Hsp90 and kinases. Clearly, our analysis of sacsin structure has raised multiple intriguing questions that will require both biochemical and structural determinations to define mechanism.

Interestingly the recombinant sacsin HEPN has itself been shown to bind nucleotides, with GTP binding at a molar ratio of one GTP per HEPN dimer. GDP and ATP were also shown to bind to the sacsin HEPN domain but at a lower affinity, 3- and 10-fold less respectively [28]. This suggests that nucleotide binding beyond the SR1 domains may influence sacsin function.

It is a common feature of chaperone machines that they are driven by ATP binding and hydrolysis to assist protein folding and unfolding [30]. Therefore, our analysis would be concordant with sacsin functioning as an ATP-driven molecular chaperone. If this is the case a key challenge will be to identify sacsin’s clients. One candidate group of proteins are intermediate filaments. Significant reorganisation of the intermediate filament cytoskeleton is observed in sacsin null cells [11,13,14,16,17]. This includes the formation of perinuclear accumulations of vimentin in sacsin knockout SH-SY5Y cells and ARSACS patient dermal fibroblasts [16], as well as abnormal bundling of non-phosphorylated neurofilament in neurons from sacsin knockout mice [17] and aggregation of glial fibrillary acidic protein in glial cells [31]. Unexpectedly, heterologous expression of isolated sacsin domains can modulate neurofilament assembly [17]. This includes the isolated UBL, SIRT 1 and J-domains, which all modified neurofilament assembly *in vivo*, with the SIRT1 and the J-domain were having opposing effects, by respectively promoting and preventing filament assembly. The intermediate filament phenotype of motor neurons from the sacsin knockout mice was also altered by expression of the sacsin SIRT1 or J-domain, with partial resolution of existing neurofilament bundles. That these isolated domains of sacsin influence neurofilament organisation is perhaps surprising but would again be consistent with the full-length protein functioning as a chaperone for intermediate filament assembly or disassembly.

Hsp90 works with other chaperones and cochaperones as part of a larger protein folding and remodelling machinery. Of particular importance is Hsp90’s collaboration with Hsp70 in protein folding and other chaperone functions [32]. The presence of the J-domain in sacsin implies that it also requires a Hsp70 partner for its function. It also seems likely that if sacsin is an ATP-driven chaperone its function could be regulated by interacting partners acting as cochaperones, as is the case for Hsp90 and other chaperones. More evidence for sacsin functioning in a chaperone network comes from its putative interactome which includes a chaperone cluster [19].

Finally, our analysis indicates that for the Hsp90 inhibitors we investigated, that target the ATP-binding site, may not bind sacsin. This is important as these drugs are in clinical trials as therapeutics for cancers and other diseases, such that off target effects on sacsin would be undesirable.

## Author Contributions

Conceptualization, J.P.C. and C.P.; Methodology, C.P. and L.E.L.R.; Formal Analysis, Methodology, C.P. and L.E.L.R.; Investigation, C.P. and L.E.L.R.; Data Curation, C.P.; Writing – Original Draft Preparation, J.P.C., C.P. and L.P.; Writing – Review & Editing, J.P.C., C.P., and L.P.; Visualization, C.P. and L.P.; Supervision, J.P.C.; Project Administration, J.P.C.; Funding Acquisition, J.P.C.

## Supplemental figure legends

**Figure S1.**
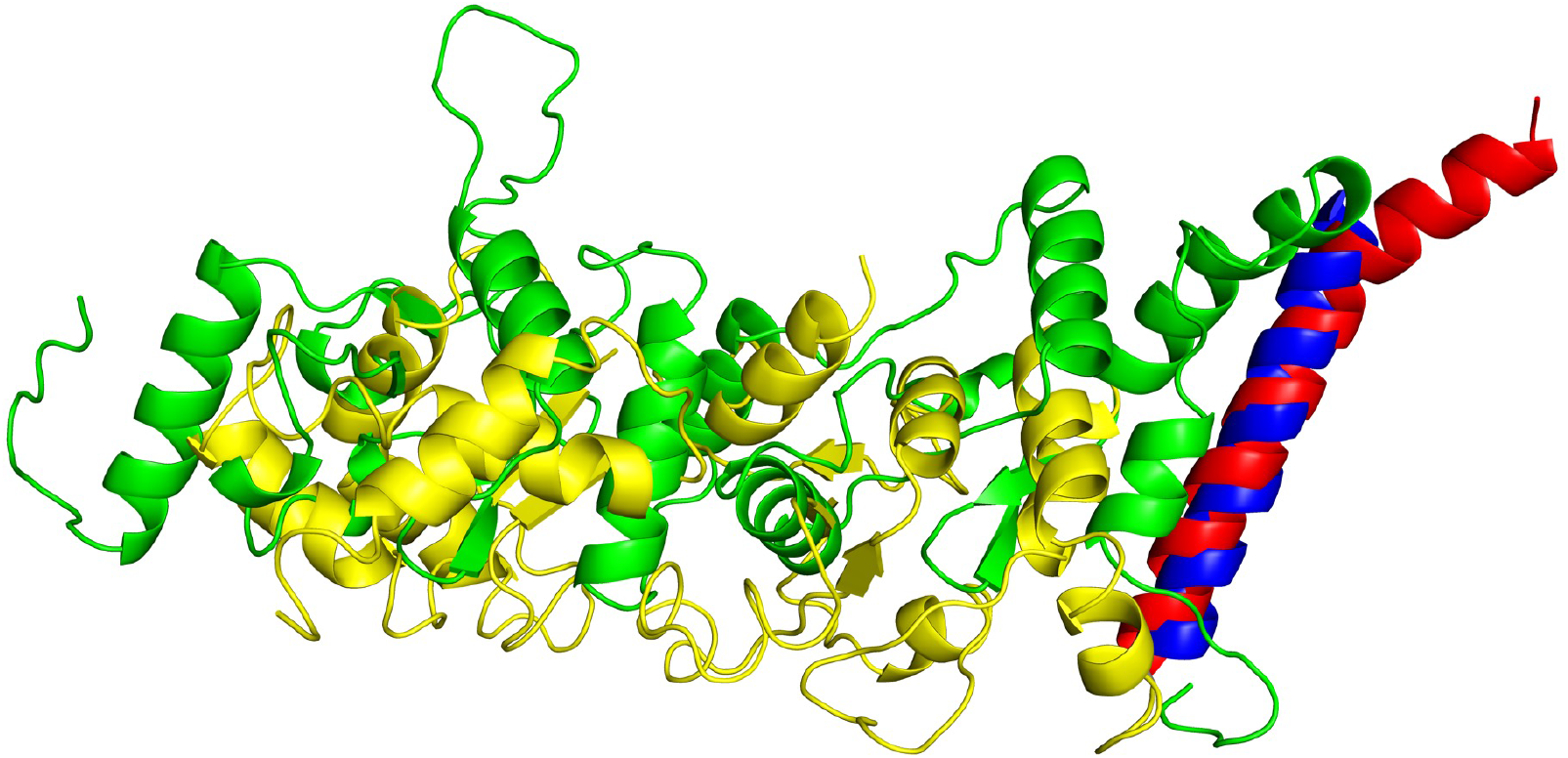
Hsp90 middle and C-terminal regions and corresponding segments of sacsin that do not superimpose. PyMol cartoon of Hsp90 (yellow and blue, residues 386 to 677) and sacsin (green and red, residues 487 to 772). The blue long helix of the Hsp90 middle domain and that of sacsin are superimposed. The rest of the structure shows no immediately recognisable common topology.

**Figure S2.**
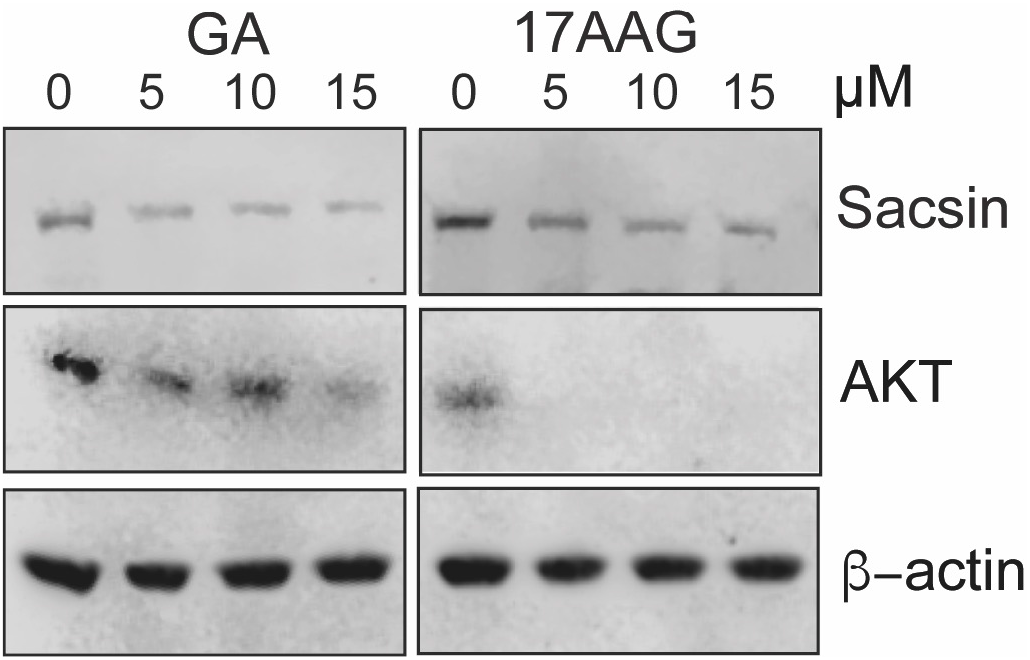
Hsp90 inhibitors reduce cellular levels of sacsin. SH-SY5Y cells were treated with the Hsp90 inhibitors geldanamycin (GA) or tanespimycin (17-AAG) (NB), and a relevant vehicle control for 24 hours. Total levels of sacsin and the known Hsp90 client protein AKT were then determined by immunoblotting. β-actin was also immunoblotted as a loading control.

